# Differentiated Oral Epithelial Cells Support the HPV Life Cycle

**DOI:** 10.1101/2023.03.08.531611

**Authors:** William T Seaman, Thatsanee Saladyanant, Victoria Madden, Jennifer Webster-Cyriaque

## Abstract

Human Papillomavirus (HPV) associated oral disease continues to increase, both in the context of immune competence and of immune suppression. There are few models of oral HPV infection and current models are laborious. We hypothesized that differentiated oral epithelial cells could support the HPV life cycle. Clinical HPV16 cloned episomes were introduced into differentiated oral epithelial cells (OKF6tert1). Viral and cellular gene expression was assessed in the presence or absence of sodium butyrate, a differentiating agent that moved the cells to full terminal differentiation. Detection of keratin 10, cross-linked involucrin, and loricrin in the presence and absence of sodium butyrate confirmed terminal differentiation. Increasing sodium butyrate concentrations in the absence of HPV, were associated with decreased suprabasal markers and increased terminal differentiation markers. However, in the presence of HPV and of increasing sodium butyrate concentrations, both mitotic and suprabasal markers were increased and the terminal differentiation marker, loricrin, decreased. In this unique differentiated state, early and late viral gene products were detected including spliced mRNAs for E6*, E1^E4, and L1. E7 and L1 proteins were detected. The ratio of late (E1^E4) to early (E6/E7) transcripts in HPV16+ OKF6tert1 cells was distinct compared to HPV16+ C33a cells. Consistent with permissive HPV replication, DNA damage responses (phospho-chk2, gamma-H2AX), HPV E2-dependent LCR transactivation, and DNase-resistant particles were detected and visualized by transmission electron microscopy. In sum, monolayers of differentiated immortalized oral epithelial cells supported the full HPV life cycle. HPV may optimize the differentiation state of oral epithelial cells to facilitate its replication.

## Introduction

The link between HPV and cervical cancer has been well established (3, 44). Less information exists for the role HPV plays in the establishment of oral cancers. Recent data from our lab and others has shown that HPV is associated with over 25% of oral cancers (1, 5, 17, 39, 46, 53). High-risk types of HPV DNA have been consistently detected in 20% of head and neck squamous cell carcinomas (HNSCC) overall with most originating in the oropharynx. HPV presence in tonsillar cancers and base of tongue cancers suggests that similar to its role in cervical transformation, HPV may transform oral epithelium as well. There is mounting evidence that oral cancers are on the rise. The incidence of tonsillar cancer, a subset of head and neck squamous cell carcinoma (HNSCC), increased 2% to 3% annually from 1973 to 2001 (48). We have shown that the cancers that develop in a subset of younger non-smoking/non-drinking (NS/ND) patients were 70% HPV positive (1).

HPV requires the differentiation program of keratinocytes to carry out its lifecyle (59). There are numerous cellular factors that are tied to differentiation that have been shown to be important to the HPV life cycle. Differentiation associated transcription factors NF1, AP1, OCT1, CEBP, SP1, YY1 and KRF-1 are critical to modulation of HPV promoters. Activation of DNA damage responses are manipulated by HPV for regulation of the differentiation dependent life cycle (36). KLF13 has recently been shown to be critical for genome amplification and viral gene expression as it modulates STAT5 and ATM DNA damage responses (56). RNA binding proteins and splicing factors that are differentiation associated are important to the proper splicing of several HPV transcripts including E6* and L1 (24). The host factors above, among others, exemplify the importance of differentiation to the HPV life cycle.

Studying the viral life cycle of HPV in oral epithelia has been hampered by the absence of a suitable tissue culture system. Most culture systems used to study the HPV lifecycle have relied on primary keratinocytes grown as raft cultures to recapitulate the epithelial differentiation with terminal differentiation associated with late gene expression and virion production (34). While valuable information concerning HPV has been obtained using these culture systems they are labor intensive and require a reliable supply of tissue to produce primary cultures. More recently primary keratinocyte culture monolayers that can be induced to differentiate by changing growth conditions with calcium or methylcellulose have been used to obtain productive HPV infection in culture (35, 43). This method also relies on an adequate supply of primary keratinocyte cultures. An additional problem that exists is the introduction of HPV into the cells. Obtaining HPV inoculum suitable for infecting cells has been arduous given the tissue culture requirements necessary to generate a productive viral infection. Typically, HPV genomes are released from a plasmid clone by restriction digest followed by electroporation of linearized genomes and recircularization within cultured cells to generate episomes (34). Adenoviral vectors containing the HPV16 genome flanked by loxp sequences have been used to introduce HPV16 episomes into primary cervical keratinocytes by transduction followed by Cre/lox recombination (32). More recently, a plasmid-based system utilizing organotypic cultures in conjunction with Cre/lox recombination for the introduction of HPV18 episomes has been described (54).

Identifying the role that HPV plays in the infection of oral epithelial cells is necessary to understanding the development of oral tumors. We have developed a simple culture system utilizing full length wild type oral HPV clinical isolates and immortalized oral keratinocytes to investigate the HPV lifecycle and facilitate the definition of viral and cellular factors critical to the spread of HPV in the oral cavity. We have built upon the previously described systems to generate loxP-containing HPV clones from oral clinical samples by PCR amplifying entire HPV genomes with type-specific primers that include loxP sequences. An HPV16 genomic clone flanked by loxP sites was then used to introduce episomes into keratinocytes. In this study, these HPV16 episomes were introduced into the previously described telomerase-immortalized, oral keratinocyte cell line, OKF6tert1 (7). Introduction of episomes into these cells resulted in the replication of episomes. Transfected cells expressed early and late viral mRNAs and effected cellular responses similar to other culture systems. In these studies sodium butyrate (NaB), an inhibitor of class I and II HDACs and inducer of cellular senescence was used as a differentiating agent (16). Using this culture system, HPV viral particles with packaged episomes were produced indicating that these cells were capable of supporting the full viral life cycle.

## Materials and Methods

### Cell culture

OKF6tert1 cells are a telomerase immortalized oral keratinocyte cell line and were provided by James Rheinwald and propagated as described (7). Briefly, OKF6tert1 cells were grown in Keratinocyte-SF (KSF) media (Gibco) supplemented with 0.2 ng/ml EGF and 25 μg/ml BPE. CaCl_2_ was added to a final concentration of 0.4 mM. C33a cells are a HPV(-) cervical carcinoma cell line and were grown in DMEM containing 10% fetal bovine serum (FBS). Caski cells were grown in Dulbecco’s modified Eagles medium supplemented with 10% FBS. All cells were maintained in a humidified incubator at 37° C with 5% CO_2_. Cells were transfected with Fugene-6 (Promega) according to the manufacturer’s instructions. Prior to treatment with NaB or CaCl2, cells were washed once with PBS and overlaid with fresh KSF media without supplements. After 24 hours, media was removed and replaced with KSF media without supplements containing either increasing concentrations of NaB or 2.0 mM CaCl2.

### Whole genome amplification

Isolation of genomic DNA from oral cancer biopsy has been previously described (46). Whole genome amplification of HPV16 from oral cancer DNA was performed with an Expand High Fidelity PCR system (Roche) according to the manufacturer’s instructions using HPV type 16-specific primers, 16loxpF and 16loxpR. One nanogram of genomic DNA was used in the reaction. Thermal cycle conditions for amplification were as follows: 94°C 2 minute initial denaturation followed by 10 cycles of 94°C for 15 seconds (denature), 60°C for 30 seconds (primer annealing) and 68°C for 7 minutes (extension). These 10 cycles were followed 20 cycles of the same thermal cycle profile except that 15 seconds were added to each successive 7 minute extension at 68°C. PCR products were analyzed by agarose gel electrophoresis.

### Plasmids and primers

The Cre expression plasmid, pCre-CMVPro, was a gift from the UNC Gene Therapy Core Facility. Primer sequences used for PCR amplification and shown in Table S1. All primers were manufactured at the UNC Nucleic Acid Core Facility. HPV16 whole genome PCR product was cloned into the HindIII/XmaI sites of pcDNA3 (Invitrogen) to construct pHPV16loxp. HPV16 E6 and E7 expression plasmids, pcDNAHPV16E6 and pcDNAHPV16E7 were constructed by PCR amplification of either the E6 coding region (83–559) with HPV16E6F/R primers or the E7 coding region (562–858) with HPV16E7F/R primers using Caski cell DNA as a template followed by cloning into the HindIII/XhoI sites of pcDNA3. HPV16 E2 expression plasmid, pcDNAHPV16E2 was constructed by PCR amplification of the E2 coding region (2755–3852) with HPV16E2F/R primers using Caski cell DNA as a template followed by cloning into the HindIII/XhoI sites of pcDNA3. pSVGFPloxp was constructed by inserting GFP under the control of the SV40 promoter flanked by loxp sites into pcDNA3Primer sequences for PCR amplification are shown in Supplemental Table 1. All plasmid constructs were verified by sequencing.

**Table 1.** OKF6tert1 cells are differentiated and support HPV viral replication. **A.** Summary of OKF6tert1 cells differentiation state with and without NaB **B.** Visual and charted summary of OKF6tert1 cells containing HPV16 episomes. Differentiation state and viral gene expression were examined with and without NaB. (–) not detected (+) detected (ND) not done

A. OKF6
|  | No treatment | 0.6 mM NaB |
| --- | --- | --- |
| Monomeric involucrin | + | + |
| Crosslinked involucrin | + | +++ |
| Loricrin | + | +++ |
| phospho serine10 H3 | ++ | ++ |
| Keratin 10 | ++ | ++ (decreases with >0.6 mM) |
| Acetylated H3 | + | +++ |

| OKF6- HPV16 | No treatment | 0.6 mM NaB |
| --- | --- | --- |
| Loricrin | ++++ | ++ |
| Keratin 10 | + | ++ |
| phospho serine10 H3 | + | ++ |
| E6/E7 expression | + | ++ |
| E6* expression | + | ++ |
| E1^E4 expression | + | ++++ |
| L1 expression | + | + |
| DNase-resistant particles | + | + |

### RT-qPCR

Total RNA was isolated from transfected cells using Trizol according to the manufacturer’s instructions (Invitrogen). Briefly, cells were washed twice with 1X PBS before the addition of Trizol. Total RNA was incubated with RQ1 RNase-free DNase (Promega) for 45 minutes at 37° C prior use in reverse transcription reactions. After incubation DNase activity was destroyed by heating to 70° C for 10 minutes. 200 ng of total RNA was reverse transcribed using SuperScript III (Invitrogen) and random primers. Reactions were performed at 50° C for 1 hour. Following incubation, reactions were diluted to 100 μl with water and 2 μl were used for qPCR. Real time PCR reactions were performed in triplicate using Roche Lightcycler Sybr Green master mix. Each reaction consisted of 1X Roche Lightcycler Sybr Green master mix and gene-specific primers (Supplemental Table 1) in a 10 μl reaction. Thermal cycle conditions consisted of an initial denaturation incubation at 95° C for 10 minutes followed by 45 cycles of alternating 95° C incubations for 10 seconds, 60° C incubations for 10 seconds and 72° C incubations for 30 seconds. Fluorescence was detected after every 72° C extension incubation. GAPDH levels were used to normalize experimental gene transcript levels.

### Western blot analysis

Total protein was isolated from cells using Trizol (Invitrogen) according to the manufacturer’s instructions. Final protein pellets were resuspended in 2X sample buffer (125 mM Tris, 8.0, 4% SDS, 20% glycerol, 50 mM DTT). Pellets in buffer were incubated at 65 C until dissolved. Twenty-five micrograms of total protein were loaded onto precast NuPage 4-12% Bis-Tris gels (Invitrogen). Proteins were electroblotted to nitrocellulose paper using western blot transfer buffer (25 mM Tris base, 192 mM glycine, 20% methanol). Following transfer, blots were blocked by incubation with 5% nonfat dry milk in 1X PBS containing 0.1% Tween-20 (1X PBS-T) for 1 hour at room temperature. Following blocking, blots were incubated with primary antibody for 1 hour at room temperature. Blots were washed twice with 1X PBS-T followed by incubation for 1 hour with HRP-conjugated secondary antibody (Promega) diluted 1:10,000 in PBS-T, 5% milk. Blots were washed twice with PBS-T and protein bands were detected by ECL after incubation with SuperSignal for 5 minutes according to the manufacturer’s instructions (ThermoScientific). The following primary antibodies were used to detect proteins on western blots: anti-involucrin (sc-28557; Santa Cruz), anti-loricrin (PRB-145P; Biolegend), anti-GAPDH (sc-25778; Santa Cruz), anti-γ-H2A.X Ser 139 (sc-101696; Santa Cruz), anti-p-Chk2 Thr 68 (sc-16297-R; Santa Cruz), anti-actin (sc-1616-R; Santa Cruz), anti-p38 MAPK (9212S; Cell Signaling), anti-phospho p38MAPK (9215S; Cell Signaling), anti-p53 (sc-6243; Santa Cruz), anti-histone H3 (D1H2; Cell Signaling), anti-acetyl-histone H3 (06-599B; Millipore), anti-phospho-histone H3 (ser10) (05-598; Upstate).

### Quantitation of DNase-resistant viral DNA

Media from transfected cells were passed through a 0.45 micron filter. 200 ml of media was treated with 4 U of RQ1 RNase-free DNase (Promega) for 1 hour at 37°C. Following incubation DNase activity was destroyed by heating to 70° C for 15 minutes. DNA was isolated with a Qiagen DNeasy Blood & Tissue kit according to the manufacturer’s instructions. HPV16 DNA was detected by qPCR using 1X Roche Lightcycler Sybr Green master mix and HPV16 E7-specific primers. Thermal cycle conditions consisted of an initial denaturation incubation at 95° C for 10 minutes followed by 45 cycles of alternating 95° C incubations for 10 seconds, 60° C incubations for 10 seconds and 72° C incubations for 30 seconds. Fluorescence was detected after every 72° C extension incubation.

### Negative staining TEM

HPV particles were detected by TEM as previously described (49). Briefly, media from transfected cells was centrifuged for 20,000 X G for 30 minutes to remove cellular debris followed by centrifugation at 100,000 X G for 1 hour to pellet viral particles. Pellets were resuspended in 50 μl of water and fixed with 3% glutaraldehyde. Particles were stained with uranyl acetate and analyzed by TEM.

### Immunofluorescence Assay (IFA)

Following transfection, cells were washed with 1X PBS and fixed with methanol/acetone. Blocking was performed for 1 hour with 1X PBS containing 5% normal goat serum. For de novo infection, filtered media from transfected cells was used as an inoculum for naïve OKF6tert1. 96 hours post infection cells were fixed with methanol/acetone. HPV16 E7 was detected using mouse anti-HPV16 E7 (sc-6981; Santa Cruz Biotech) and HPV L1 was detected with CAMVIR-1 antibody (sc-47699; Santa Cruz Biotech). Fixed cells were overlaid with 1:500 dilution of primary antibody and incubated overnight at 4° C. Cells were washed three time with 1X PBS (5 minutes/wash) followed by overlaying with 1:500 dilution of Alexa Fluor 488 goat-anti mouse IgG (A28175; Pierce) for 1 hour at room temperature. To stain nuclei, cells were overlaid with DAPI for 5 minutes followed by washing three time with 1X PBS (5 minutes/wash). Fluorescence was detected with an Olympus IX81 fluorescent microscope.

## Results

### Oral epithelial cells can be induced to terminally differentiate with calcium and sodium butyrate (NaB) treatment

HPV replication is tied to epithelial differentiation (zurHausen, 2002). Both calcium and sodium butyrate (NaB) treatment have been shown to induce keratinocyte differentiation markers in epithelial cells (20, 51). To determine whether telomerase-immortalized oral keratinocytes (OKF6tert1) supported the HPV16 life cycle, differentiation status and susceptibility to calcium and sodium butyrate-induced differentiation were assessed. The detection of the keratinocyte differentiation marker, monomeric involucrin (70 kD), in the absence of CaCl_2_ or NaB treatment suggested that these cells were already differentiated (Figure 1A and 1B). Likewise, involucrin was detected by immunoflourecence in both low and high calcium conditions (data not shown/ supplementary data). At seventy-two hours post treatment with CaCl_2_ or NaB, differences in the level of monomeric involucrin were not detected by western blot analysis (Figures 1A and 1B). Involucrin cross-linking is a common occurance in the full terminal differentiation of keratinocytes. Indicative of involucrin crosslinking, additional bands at 120 kD and 240kD, were observed in cells treated with 1.5 mM CaCl_2_ (Figure 1A) and with NaB (Figure 1B). The 0.4 mM level of calcium present in the Keratinocyte serum free media (K-SFM) was not sufficient to induce the 120kD band (Figure 1A, CaCl2 (-) lane). The differentiation marker, loricrin, has been characterized as a terminal differentitation marker in normal epithelia (21) and is dysregulated in several disease states (23). Exposure of cultured keratinocytes terminal differentiating agents (CaCl_2_ or retinoic acid) upregulated loricrin mRNA and protein (22). In this study, low levels of loricrin were detected in untreated OKF6tert1 cells. Exposure of cells to increasing NaB concentrations (0.13, 0.25, 0.50, 1.0, and 2.0 mM NaB) resulted in a graduated increase in loricrin protein levels as determined by western blot analysis (Figure 1B). In contrast to loricrin, the suprabasal differentation maker Keratin 10 (K10) was detected at higher levels in the absence of NaB and at lower NaB concentration (0.13mM) and decreased with increasing NaB concentrations (0.13, 0.25, 0.50, 1.0, and 2.0 mM NaB) as determined by immunoblot (Figure 1B). Consistent with its activity as a histone deacytylase inhibitor, increasing levels of acetylated histone 3 (AcH3) were detected with increasing NaB concentrations (Figure 1B). Total histone 3 and GAPDH served as loading controls. Taken together these results suggested that at baseline, OKF6tert1 cells were in a differentiated state but could be further induced using either NaB or CaCl2 to express markers associated with full terminal differentation.

**Figure 1.**
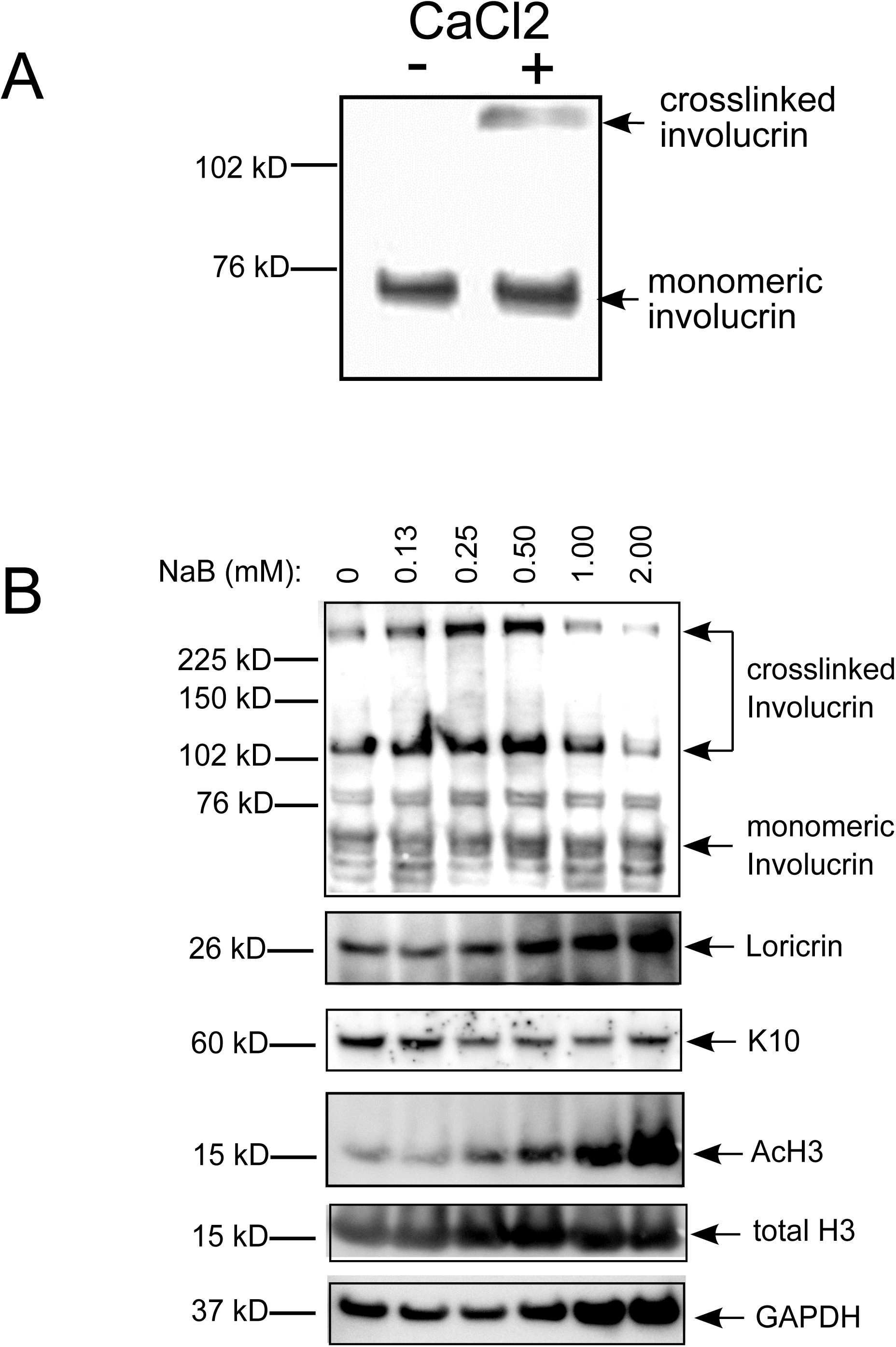
Terminal keratinocyte differentiation markers can be induced in the differentiated, immortalized keratinocyte cell line, OKF6tert1. **A.** Western blot analysis of monomeric (early differentiation) and crosslinked (late differentiation) involucrin in whole cell extracts from OKF6tert1 cells grown K-SFM containing 0.4 mM CaCl_2_ or following addition of CaCl_2_ to a final concentration of 2 mM to culture medium. **B.** Cell lysates from OKF6tert1 cells treated with vehicle or NaB for 72 hours were subjected to SDS_PAGE and western blot analysis. 0.13, 0.25, 0.50, 1 and 2 mM concentrations of NaB were assessed. Monomeric involucrin was detected in all extracts and is indicated by the arrow. A 120 kD and 250kD bands representing crosslinked involucrin were detected when cells are grown in the presence of NaB as indicated by the arrowhead. Increasing amounts of the terminal differentiation marker, loricrin, was detected with increasing concentrations of NaB. Expression of the spinous differentiation marker keratin 10 (K10) and of acetylated histone 3 (AcH3) were determined by western blot analysis. GAPDH and total H3 served as loading controls.

### Oral Episomal HPV was cloned by Cre/lox recombination and replicated in oral epithelial cells

Frequently oral and cervical cancers contain a mixture of integrated HPV16 DNA and of viralepisomes (4, 29). In some cancers, HPV16 integrates in tandem repeats of the viral genome without loss in viral genetic information (25). Thus, full length HPV16 genomes can be obtained from HPV-associated cancers if episomes or tandem repeats are present. In a previous study, we identified oral cancer biopsies containing HPV16 DNA (46). This DNA was used as a template for whole genome amplification of HPV16. Two HPV16-specific oligonucleotides were designed that primed DNA synthesis in opposite direction around the HPV16 genome (Figure 2A). The forward primer began at nucleotide 7271 immediately 5’ to the LCR. The reverse primer began at nucleotide 7270 and was immediately 3’ to the late polyA signal. The 5’ end of each primer was designed to include loxP recombination sites as well as a unique restricition enzyme recognition site. A product of ∼7.9 kb that included a 34 bp loxp sequence was PCR amplified using template DNA obtained from 4 oral cancer biopsies (D12, C9, C10 and F3) (46) (Figure 2B). Sequencing of the full length D12 clone verified a full length HPV16 genome with all of the open reading frames intact including those that encoded L1 and L2 (data not shown), suggesting that the HPV16 genome in these oral cancers existed as either a mixture of episomes and integrated DNA or as integrated tandem repeats of full length genomes. OKF6tert1 cells were cotransfected with the Cre expression vector, pCRE-CMVPro, and pHPV16loxp to determine if HPV16 episomes were generated by Cre recombination (Figure 2C). HPV16-specific qPCR assays that targeted the loxp junction were performed on total genomic DNA isolated from the OKF6-HPV cells at 24, 48, 72 and 96 hours post-cotransfection (Figure 2D). PCR primers to pcDNA (colE1F/R), independent of Cre recombinase activity, were used for qPCR copy number determination in cotransfected cells. Throughout the time course, plasmid specific colE1F/R primers detected no change in DNA copy number, reflecting the stablility of these DNA sequences in OKF6tert1 cells. A recombination event utilizing the loxp sequences flanking the HPV16 genome was expected after expression of Cre recombinase. This brought the primer binding sites for HPV16jncF/R primer pair in close proximity and allowed for PCR amplification across the loxp recombination junction of the newly generated episome (Figure 2A and 2C). Between 24 and 72 hours post transfection the episome-dependent primers detected a one log increase in the copy number of new episomes while the level of plasmid DNA, as indicated by colE1 amplification, remained constant (Figure 2D, a representative experiment is shown). Caski, containing tandem repeats of HPV16, served as a positive control. The detection of an HPV episomal PCR product was visualized on an agarose gel. The slight increase in size relative to Caski, was likely due to the insertion of the 34bp loxP sequence (Figure 2E).

**Figure 2.**
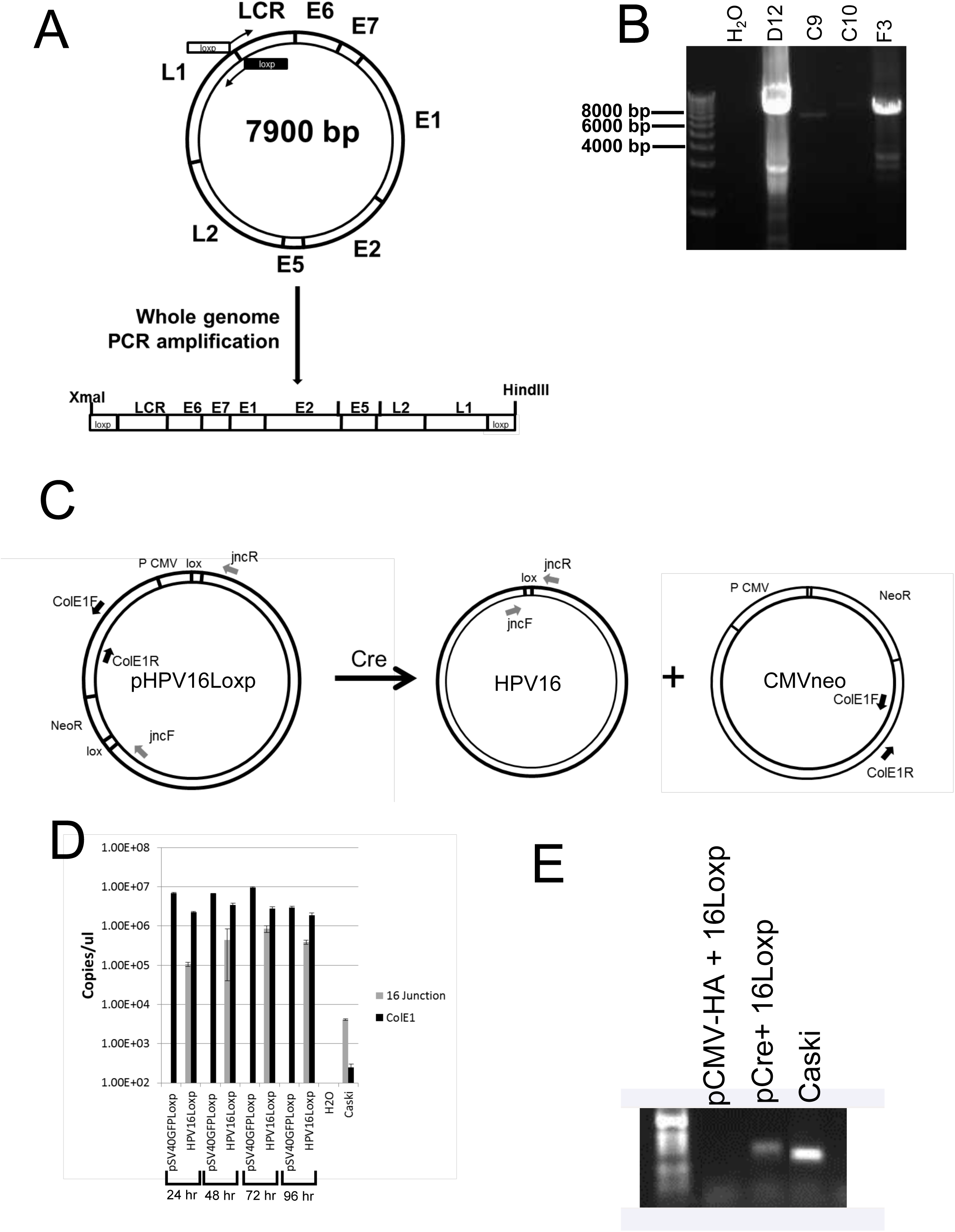
qPCR analysis of HPV16 DNA in OKF6tert1 transfected cells. **A.** Cloning strategy to generate HPV16 genomes flanked by loxp sequences. HPV16 PCR primers containing loxp sequences were used to PCR amplify the entire HPV16 genome. The direction of strand synthesis by each primer is indicated by the arrow. The boxes associated with each primer indicate the loxp sequence. The circle represents the HPV16 episome and the organization of the viral genome. The bottom line represents the linear HPV16 genome obtained by amplification with the primers. HindIII and XmaI restriction sites in the original primer sequences were used in the subsequent cloning of PCR-generated HPV16 genomes. **B.** Agarose gel analysis of whole HPV16 genomes generated by PCR. Viral genomes were PCR amplified using total DNA obtained from four HPV16-positive oral cancer biopsies. A 7.9 kbp PCR product was detected in PCR reactions containing oral cancer biopsy DNA. The resulting PCR products were cloned into pcDNA3 using the HindIII and XmaI restriction sites. The resulting plasmid, pHPV16loxp has the HPV16 genome flanked by loxp sequences between the CMV promoter and the neomycin resistance gene. **C.** Schematic representation showing the position of primers and binding sites before and after Cre/lox recombination to generate HPV16 episomes in OKF6tert1 cells. Cre/loxp recombination generates an HPV episome and another circular DNA that juxtaposes the CMV promoter next to the neomycin resistance gene. Black arrows show the position of primers that amplify the ColE1 origin of replication present in the plasmid backbone. Gray primers indicate HPV16-specific primers that are put in close proximity following recombination. **D.** DNA isolated from transfected cells was used to PCR amplify across the junction formed by Cre/loxp recombination. qPCR results for each primer pair indicated that HPV16 genomes were present in the transfected cells. Amplification with primers that cross the loxp junction indicates that episomes were increasing over time suggesting that replication of the viral genome was occurring. The gel shows the PCR product generated with primers that cross the loxp junction. The slight increase in band size relative to the Caski DNA control reflects the additional 34 bp loxp sequence.

### Markers of cellular differentiation were assessed in the context of HPV

During normal epithelial differentiation a known inverse relationship exists between expression of suprabasal markers such as K1 and K10 and terminal differentiation markers that include loricrin and fillagrin (15, 37). Differentiation markers were assessed within oral keratinocytes in the context of increasing NaB levels (Figure 3). Consistant with HDAC inhibition activity, increasing levels of AcH3 were detected with increasing concentrations of NaB (0.13, 0.25, 0.5 and 1mM) in OKF6tert1 transfected cells both in the presence or absence of HPV. Presence of HPV was confirmed by consistent detection of HPV DNA (Figure 2A). Lysates analyzed for protein expression came from the 72 hour time point. Expression of HPV in the absence of NaB increased cell differentiation in HPV transfected OKF6tert1-cells compared to SVGFP transfected cells. In the HPV expressing cells, increased expression of the terminal differentiation marker, loricrin, and decreased expression of the suprabasal marker, K10. In the presence of HPV16 the marker of mitotic activity, phosphoserine10 histone 3 (pS10-H3), was suppressed relative to vector alone. pS10-H3 is also reduced in assocation with DNA damage, hence its suppression in the presence of HPV may be associated with HPVE7 mediated DNA damage response (36). However, in the presence of increasing levels of the differentiating agent, HPV16 was associated with decreased loricrin levels and increased levels of K10. Addition of NaB increased in HPV16-associated pS10-H3 levels. In contrast, pS10-H3 levels did not appear to change with NaB treament in the absence of HPV (Figure 3B). In the presence of HPV16, modified expression of differentiation and mitotic activation markers occurred. HPV16 modulated cellular differentation processes as indicated by the markers assayed above may reflect a unique differentiation phenotype that may provide a significant advantage to the HPV lifecycle.

**Figure 3.**
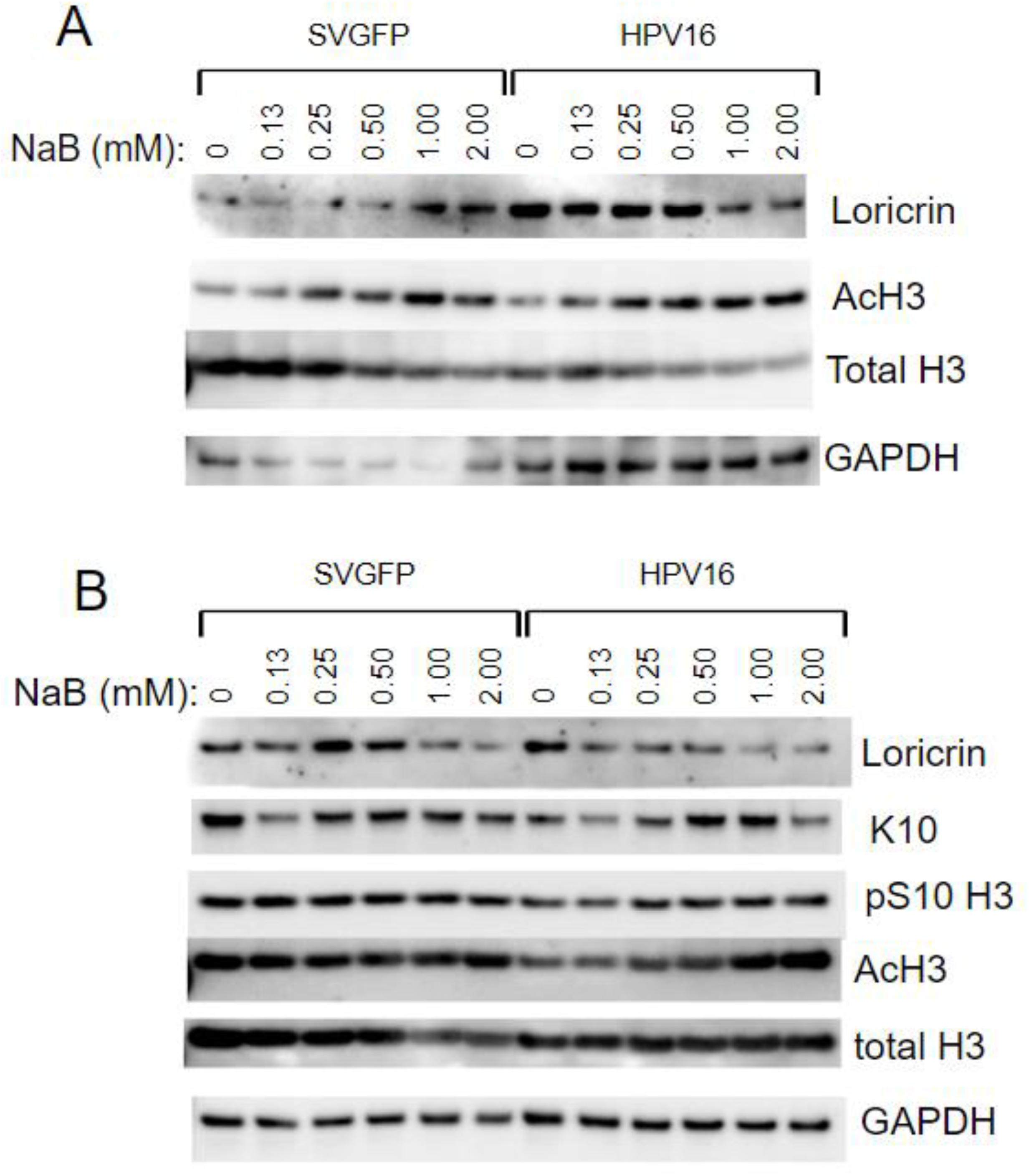
Expression of differentiation markers in OKF6tert1 cells is modified by the presence of HPV. **A.** Cells were transfected with pSVGFPloxp or pHPV16loxp. Cell lysates from OKF6tert1 cells treated with vehicle or increasing concentrations of NaB (0.13, 0.25, 0.50, 1.00 or 2.00 mM) for 72 hours were subjected to SDS-PAGE and western blot analysis. Monomeric involucrin was detected in all extracts and is indicated by the arrow. In cells transfected with pSVGFPloxp, a 120 kD band representing crosslinked involucrin was detected when cells are grown in the presence of NaB as indicated by the arrowhead. In cells transfected with pHPV16loxp, lower levels of the 120 kD band representing crosslinked involucrin were detected when cells are grown in the presence and absence of NaB as indicated by the arrowhead. Fold increase in crosslinked involucrin relative to monomeric involucrin is indicated in the graph. **B.** Cells were transfected with pSVGFPloxp or pHPV16loxp. Cell lysates from OKF6tert1 cells treated with vehicle or 0. 0.25, 0.5 or 1 mM NaB for 72 hours were subjected to SDS_PAGE and western blot analysis. Levels of cytokeratin K10, loricrin, and acetylated H3 were detected in the extracts.

### HPV early and late genes are expressed in oral epithelial cells

The study of the HPV16 lifecycle in OKF6tert1 oral keratinocytes is dependent on the expression and function of HPV16-specific genes. HPV16 late gene expression is relies on keratinocyte terminal differentiation (30, 45, 58). The effect of OKF6tert1 terminal differentiation on HPV16 gene expression was assessed. Induction of full terminal keratinocyte differentation in transfected cells was accomplished with 0.6mM NaB treatment. RT-qPCR was employed to determine whether HPV16 transcripts were expressed from episomes generated in differentiated oral keratinocytes, post-transfection.

It has been proposed that the production of E7 protein is dependent on the removal of an intron from the E6 coding region resulting in the expression of E6* (42, 52). Both in the differentiated state (0 mM NaB) and upon further differentiation of cells with 0.6 mM NaB, E6* spliced transcripts were increased as shown by RT-PCR (Figure 4A, representative experiment, three biologic replicates have been performed). With the addition of NaB, HPVE6* transcripts were enhanced by 1.4 fold as determined by RT-qPCR (Figure 4D). This spliced transcript was the major early mRNA detected in OKF6tert1cells and E6* expression has been described in HPV16-associated neoplasia and cancer (2, 9, 47, 57). Western blot analysis for E7 protein detected a 17 kD band in HPV16-containing cells regardless of NaB status (Figure 4C). However, RT-qPCR detected an increase in E6/E7 transcripts with increasing NaB as reflected by a decrease in the ΔCt values (Figure 4B). Amplification of the E6/E7 and E6* mRNAs suggested that early viral RNAs were expressed from these episomes. Transcripts containing E6/E7 sequences were detected by 48 hours, peaked after 72 hours and began to decrease at 96 hours post transfection (Figure 4E). A similar trend was detected for expression of the late E1^E4 transcript (Figure 4E).

**Figure 4.**
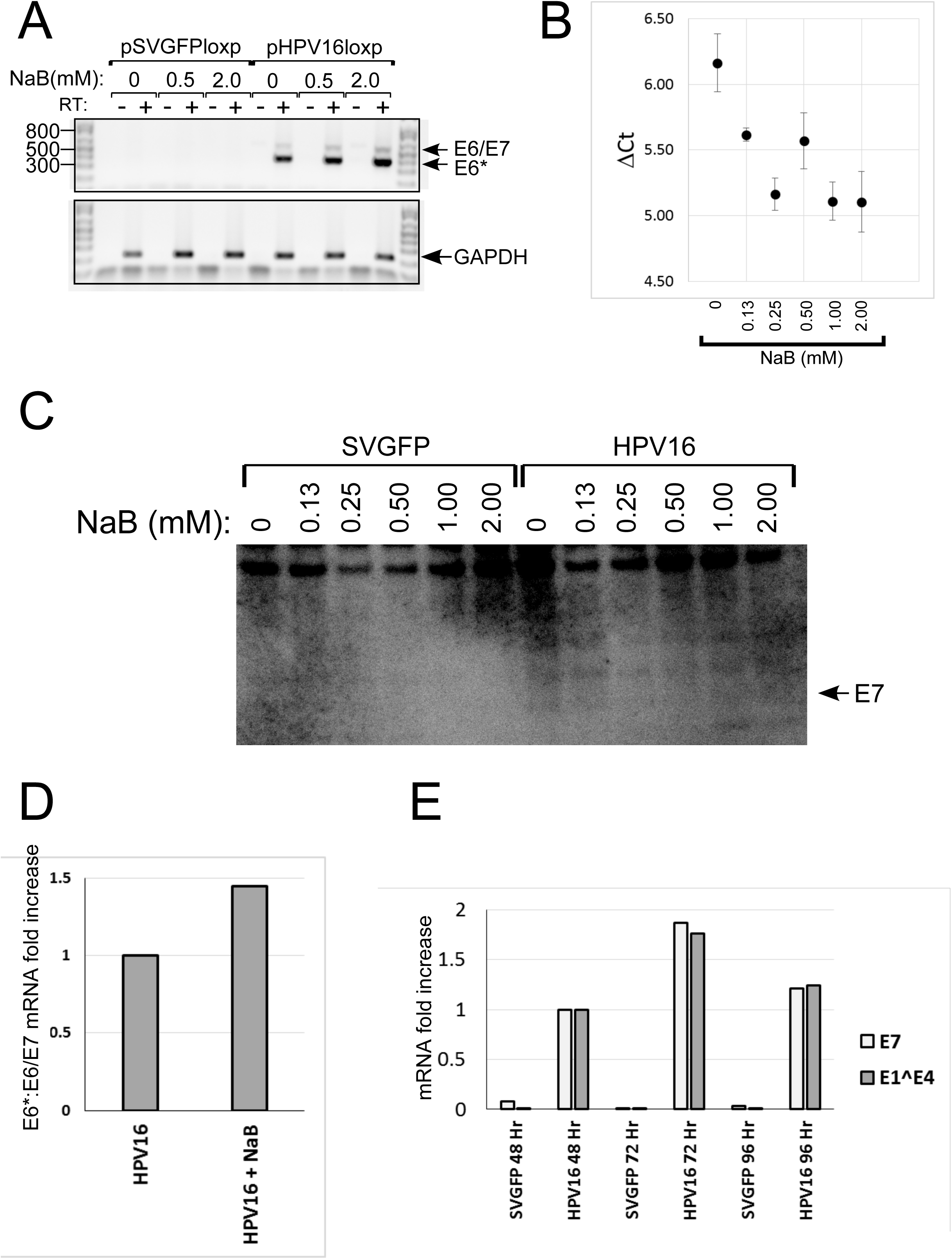
Sodium butyrate increases HPV16 early gene products in transfected OKF6tert1 cells. **A.** E6* is the major transcripts detected in OKF6tert1 cells harboring HPV16 episomes. Total RNA was isolated from pHPV16loxp-transfected OKF6tert1 cells 72 after treatment with either 0.0, 0.5 or 2.0 mM NaB. Reverse transcription using random primers and Superscript III was performed on total RNA. PCR was performed using HPVE6/E7 primers. Bands representing E6/E7 (unspliced) and E6* (spliced) transcripts are indicated. **B.** HPV16 primers specific for either E6/E7 or E6* transcripts were used to detect HPV16 cDNAs using Sybrgreen-based qPCR. The ratio of E6* to E6/E7 transcripts was determined after normalization of both signals to GAPDH. **C.** E7 protein is made in OKF6tert1 cells harboring HPV16 episomes. Total protein extracts from transfected OKF6tert1 cells were subjected to SDS-PAGE and western blot analysis using mouse anti-HPV16 E7 antibody. A 17 kD band was detected in cells containing HPV16 episomes. **D.** Treatment of OKF6tert1 cells with NaB results in increased splicing of E6/E7 mRNA to produce E6* transcripts. RNA from transfected OKF6tert1 cells was used in RT-qPCR. **E.** Total RNA was isolated from OKF6tert1 cells 48, 72 and 96 hours after cotransfection with pCre and either pSVGFPloxp (vector) or pHPV16loxp (HPV16) and used for RT-qPCR. RT-qPCR was used to detect early (HPV16 E7) and late (E1^E4) gene expression in OKF6tert1 cells. The fold increase was determined using GAPDH levels as a calibrator.

The E1^E4 protein is encoded by a mRNA that utilizes a major splice donor site at nt880 (SD880) and a major splice acceptor site at nt3358 (SA3358) (50). To detect the class of HPV late transcripts, primers specific for HPV16 spliced E1^E4 were used. RT-PCR using E1^E4-specific primers and RNA obtained from OKF6tert1 cells harboring HPV16 episomes indicated that SD880 and SA3358 were being utilized to create the late E1^E4 transcript. Both in the differentiated state (0 mM NaB) and upon further differentiation of cells with 0.5 and 2.0 mM NaB concentrations, E1^E4 spliced transcripts were detected as shown by visualization of a 100bp RT-PCR product on an agarose gel (Figure 5A). Quantitative RT-PCR detected a 4 fold increase in E1^E4 specific mRNA in HPV16 transfected cells upon NaB exposure, over and above basal E1^E4 expression levels in HPV16 transfected cells that were untreated, (p=0.028) (Figure 5B). E1^E4 expression was compared in oral and cervical epithelial cells, OKF6tert1 and C33a respectively. While NaB resulted in a 50% increase in expression of E1^E4 in OKF6tert1 cells, little change was detected in C33a cells treated with NaB (Figure 5C, representative experiment shown). The ratio of late (E1^E4) to early (E7) transcripts were compared in differentiated OKF6tert1 cells and in undifferentiated C33a cervical cancer cells. The OKF6tert1 cells demonstrated a two fold higher ratio of late to early transcripts compared to the C33a cells, although this difference was not statistically significant (p>0.05) (Figure 5D). RT-qPCR detected an increase in E1^E4 transcripts with increasing NaB as reflected by a decrease in the ΔCt values (Figure 5E, left panel). Likewise, RT-qPCR detected a decrease in spliced L1 transcripts with increasing NaB as reflected by a increase in the ΔCt values (Figure 5E, right panel). These results suggest that E1^E4 and L1 have distinct differentiation dependant requirements for their expression. These studies were carried out at with at least three bioogical replicates with each replicate having technical triplicates performed. L1 protein was detected by immunofluoresence utilizing Alexafluor 488 in HPV-containing OKF6tert1 cells. Perinuclear staining (green) was observed for L1 and was increased after treatment with NaB (Figure 5F). Nuclei were Dapi stained (blue). In sum, these data indicated that the level of keratinocyte differentiation was sufficient for HPV16 early and late gene expression in this oral epithelial cell culture system.

**Figure 5.**
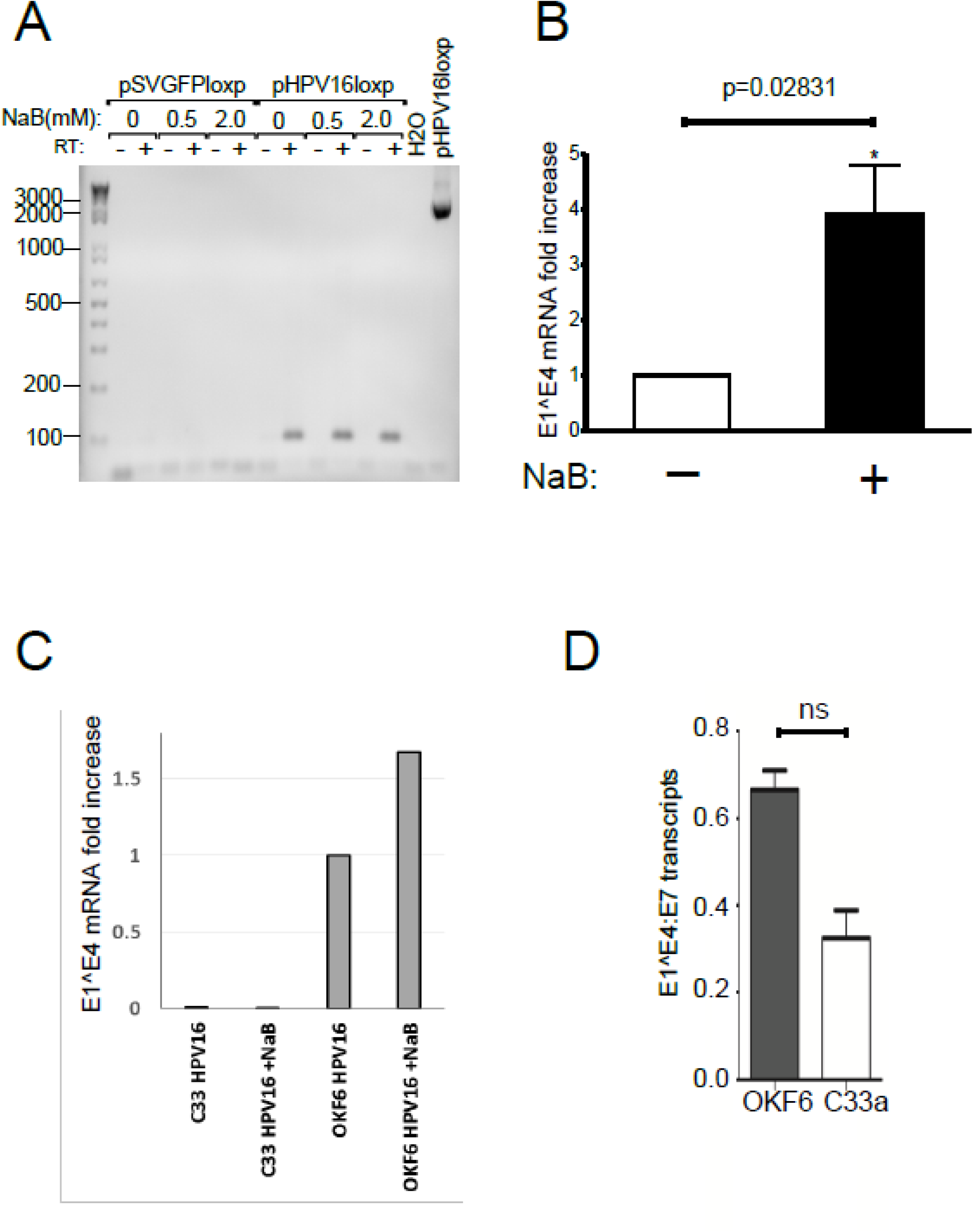
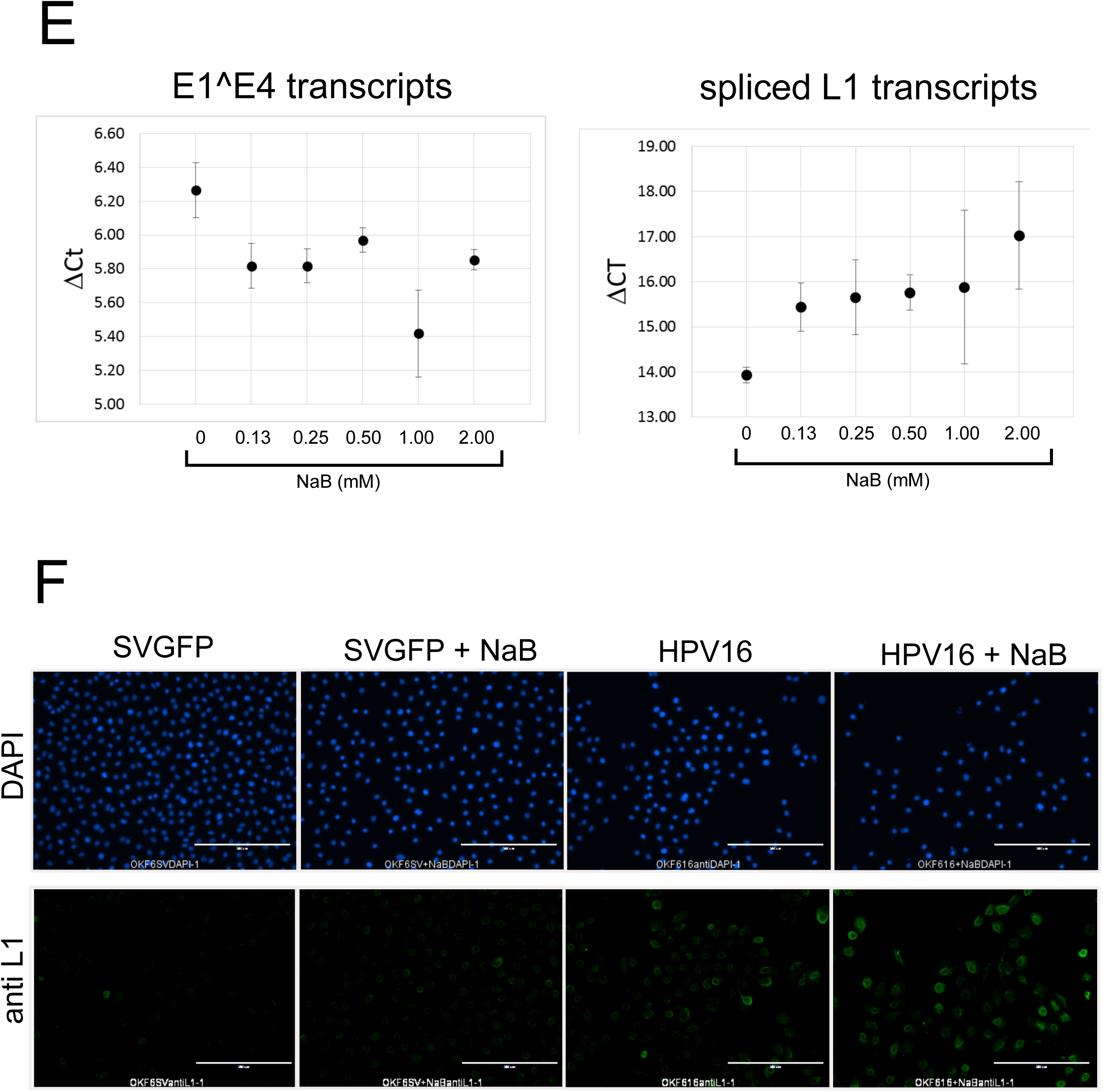
HPV16 late gene products can be detected in transfected OKF6tert1 cells. **A.** Total RNA was isolated from OKF6tert1 cells 48, 72 and 96 hours after cotransfection with pCre and either pSVGFPloxp (vector) or pHPV16loxp (HPV16) and used for RT-qPCR. Reverse transcription using random primers and Superscript III was performed on total RNA. HPV16 primers specific for spliced E1^E4 transcripts were used to detect HPV16 cDNAs using Sybrgreen-based qPCR. Fold increases were determined using GAPDH as a calibrator. **B.** RT-qPCR analysis of HPV16 E1^E4 gene expression in C33a (HPV(-) cervical carcinoma) or OKF6tert1 cells after induction of terminal differentiation with 0.6 mM NaB. The fold increase was determined using GAPDH levels as a calibrator. Statistical analysis was performed using 3 biological replicates. **C.** OKF6tert1 cells harboring HPV16 episomes express higher levels of viral late transcripts (E1^E4) than C33a cells harboring HPV16 episomes. HPV16 episomes were introduced into OKF6tert1 and C33a cells and total RNA was harvested 72 hours post transfection. RT-qPCR was performed on total RNA. **D.** Induction of keratinocyte differentiation by NaB Causes a rise in E1^E4 transcripts (decreasing ΔCt values relative to GAPDH calibrator) and a decrease in spliced L1 transcripts (increasing ΔCt values relative to GAPDH calibrator). **E.** HPV16 L1 protein was detected in OKF6tert1 cells by IFA. Following transfection of OKF6tert1 cells with either pSVGFPloxp or pHPV16loxp, cells were either treated with vehicle or 0.6 mM NaB for 72 hours. Cells were fixed with ice-cold acetone:methanol (1:1) and incubated with 1:500 mouse anti-HPV16 L1 antibody (Camvir-1). Bound anti-HPV16L1 antibody was detected with Alexfluor488 goat anti-mouse IgG (Life Technologies). Cells were stained with DAPI to visualize nuclei. **F.** HPV16 L1 and HIVtat are coexpressed in some transfected cells.

### HPV early and late genes are functional in oral epithelial cells

The function of several HPV16 proteins in OKF6tert1 cells was investigated in transient transfection experiments. It is well established that expression of HPV E7 in primary foreskin keratinocytes results in double-stranded DNA breaks and induces a DNA damage response (10, 33). OKF6tert1 keratinocytes were transfected with HPV16 E6, HPV16 E7 or both expression plasmids to determine whether these viral proteins induced a DNA damage response. E7 and E6 transfected OKF6tert1 cells revealed an increase in γ-H2AX expression, a marker of DNA damage (Figure 6A lanes 2 and 3). Likewise, introduction of HPV16 episomes into OKF6tert1 cells resulted in an increase in phospho-chk2 protein levels (Figure 6B, lanes 1 and 2). These results suggested that the DNA damage occuring in OKF6tert1 cells harboring HPV16 episomes was at least partially due to HPV gene expression (Figure 6 A and B). HPV16 E5 activates proliferative responses through mitogen activated protein kinases (18). Three variants of HPV16 E5 were overexpressed in these oral epithelial cells; an N-terminal HA tagged E5 (lane3), a C-terminal HA tagged (lane 4), or untagged (lane5). These were compared to the full length HPV16 deleted for E5 that served as a negative control (lane 2). HPV16 E5 expression in OKF6tert1 cells resulted in phosphorylation of p38 MAPK (lanes 3-5), while the HPV16 whole genome deleted for E5 demonstrated reduced levels of p-p38 (lane2) (Fig 6C). HPV16 E2 can act as a positive regulator of HPV16 gene expression by binding to the HPV16 LCR and controlling viral mRNA transcription (28). OKF6tert1 keratinocytes were cotransfected with a HPV16 E2 expression plasmid and a HPV16 LCR/luciferase reporter construct to determine whether this viral protein was functional. 2ug of HPV16 E2 plasmid significantly increased HPV LCR promoter activity as determined by an 2 fold increase in luciferase activity compared to transfected cells not expressing HPV16 E2 (p=0.0175). Doses ranging from 0.125-1 ug did not result in increased activity that was statistically significant (Figure 6D). HPV16 proteins expressed from either individual plasmids or from the HPV16 episome were functional in OKF6tert1 keratinocytes suggesting that these cells were suitable for investigating the HPV16 lifecycle.

**Figure 6.**
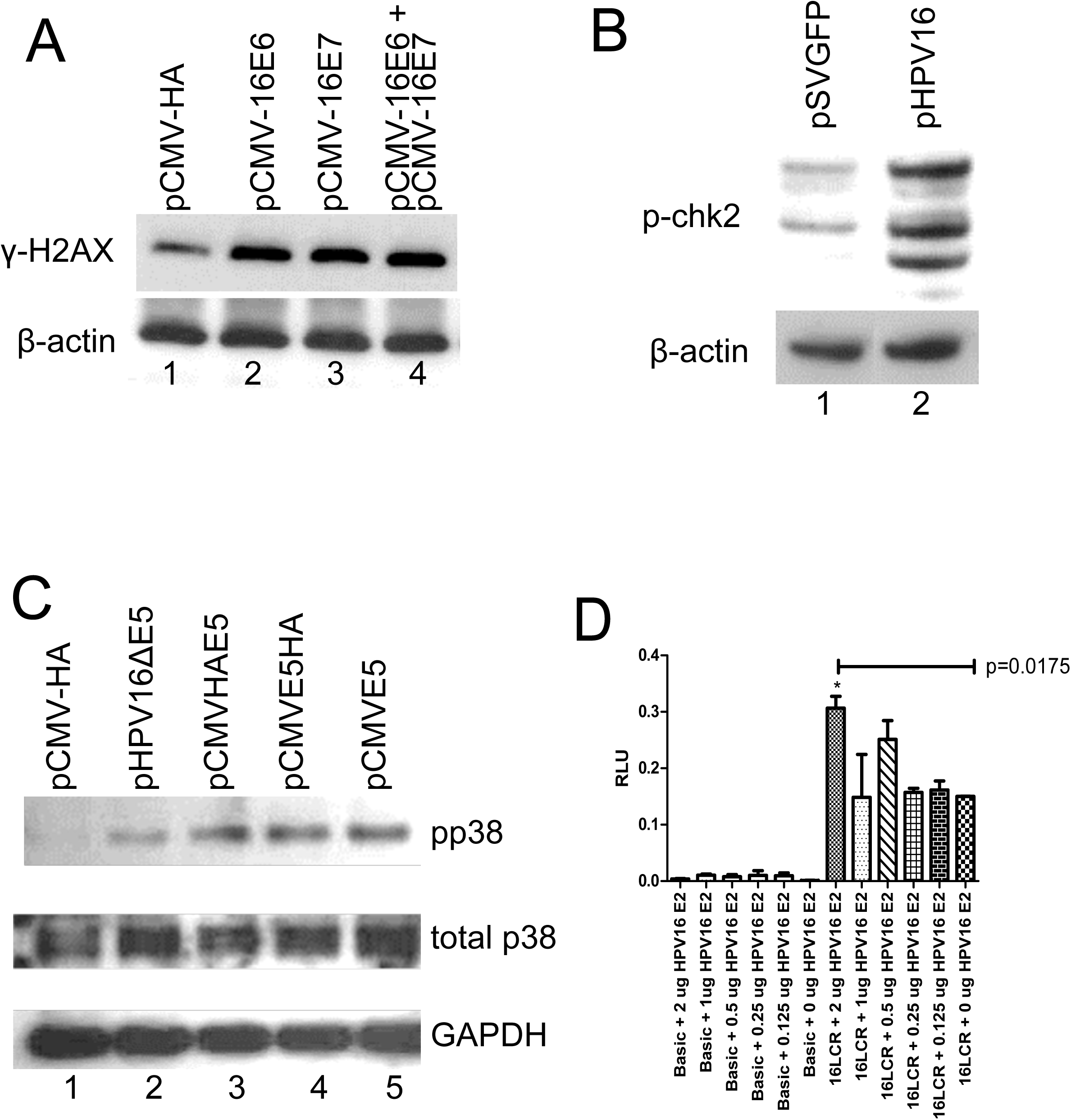
HPV16 proteins are functional in transfected OKF6tert1 cells. **A.** OKF6tert1 cells were transfected with HPV16 E6 and/or E7 expression vectors and whole cell extracts were examined by western blot analysis for the DNA damage marker, γ-H2AX. **B.** Whole cell extracts from OKF6tert1 cells harboring HPV16 episomes were analyzed for the presence of the DNA damage sensor protein phospho-chk2 by western blot analysis. **C.** HPV16 E5 expression in OKF6tert1 cells induced a cell proliferative response as determined by western blot analysis to detect phospho-p38. **D.** Expression of HPV16 E2 in OKF6tert1 cells induces luciferase expression under the control of the HPV16 LCR. E-F.

### Transfection of cloned oral HPV results in the generation of DNase resistant particles

An important aspect of the HPV16 lifecycle is the production of viral particles. The presence of DNase resistant HPV16 DNA in the supernatant media of differentiated OKF6tert1 cells haboring viral episomes was determined by qPCR. Following DNase treatment of media, DNA was isolated and HPV16 DNA levels were determined by targeting the E7 region of the genome for quantitation of DNA copy number. For each condition, media was spiked with plasmid DNA prior to DNase treatment to monitor the enzymatic removal of free DNA from the media. HPV16 DNA could not be detected above background in untransfected or control transfected OKF6tert1 cell media. HPV16 DNA was detected after the addition of plasmid DNA to the media followed by subsequent DNA isolation. Treatment of the spiked media with DNase resulted in elimination of HPV16 DNA signal, in SVGFP-containing cells, indicating that the DNase removed the free DNA from the culture media. Prior to DNase treatment, a strong HPV16 DNA signal was detected in the culture media of OKF6tert1 cells haboring HPV16 episomes. In HPV16loxp containing cells, 8×10^5^ copies of DNA were detected post DNAse treatment. The persistance of this signal in spiked media suggested that the viral episome had been packaged into particles and was not accessible to DNase degradation(Figure 7A, a representative experiment performed in triplicate is shown). HPV16 plasmid DNA and Caski DNA served as positive controls. The presence of DNase resistant HPV16 DNA in the media of NaB differentiated OKF6tert1 cells haboring viral episomes was also determined by qPCR. Since previous experiments demonstrated that NaB treatment increased late gene expression, NaB treatment was assessed for its ability to increase DNase-resistant HPV16 DNA in the culture media. Interestingly, no difference in the copy number of DNase-resistant DNA was detected in media from cells treated either with or without NaB, 5 × 10^5^ copies per ml were detected in each instance (Figure 7B).

**Figure 7.**
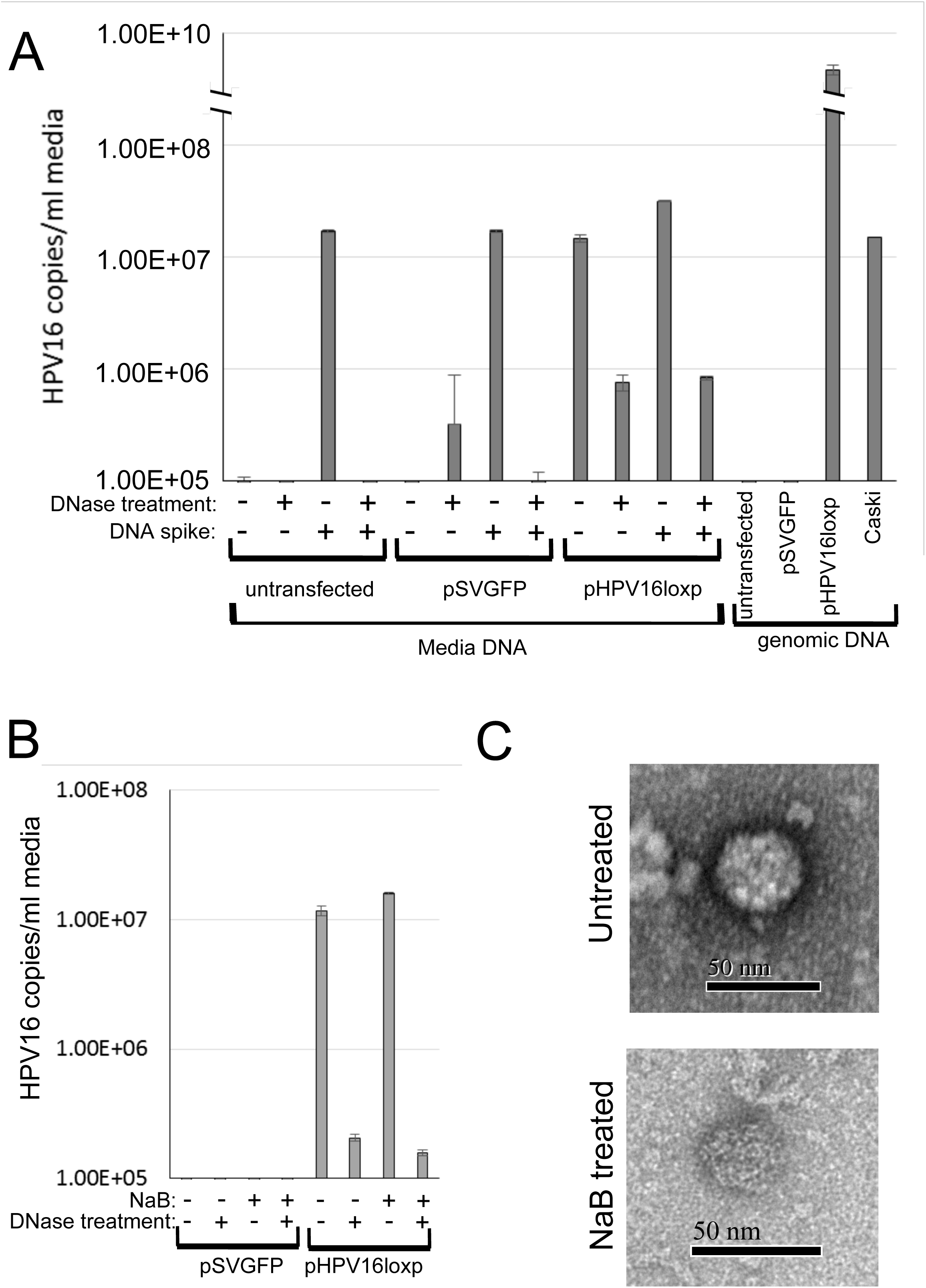
DNase-resistant HPV16 DNA present in the media of transfected OKF6tert1 cells indicates HPV16 particles are being produced by transfected cells. **A.** Media from transfected cells was filtered through a 0.45 micron filter and left untreated or treated for 1 hour with RQ1-RNase-free DNase (Promega). DNA from media was isolated using a Qiagen DNeasy kit. Four microliters of eluted DNA was used to detect HPV16 E7 qPCR. In some cases media was spiked with pHPV16loxp prior to DNA purification to monitor the effectiveness of DNase digestion. Media from pHPV16loxp cells contained HPV16 DNA that was consistently DNase-resistant suggesting that encapsulated DNA was shed by transfected OKF6tert1 cells. Media from either untransfected cells (No DNA) or pCMVNeo-transfected cells as well as purified pCMVNeo were used as a negative control. Caski genomic DNA and purified pHPV16loxp were used as positive controls. **B.** Induction of terminal differentiation in OKF6tert1 cells by 0.6 mM NaB did not significantly affect the production of DNase-resistant encapsulated HPV16 DNA as determined by qPCR. Media from cells was treated as in A after treatment with 0.6 mM NaB. HPV copy number was determined using HPV16 E7 specific primers. **C.** Transmission electron microscopy was performed on media from OKF6tert1 cells harboring HPV16 episomes. Particles ∼ 37.5 nM in size were detected in media.

The presence of viral particles in the media was also determined by transmission electron microscopy. Structures reminiscent of HPV virus like particles (VLPs) (27) were identified in the media of OKF6tert1 cells harboring HPV16 episomes (Figure 7C). A range of virion sizes have been described for HPV, ranging from 40-60nm (27). The DNase resistant particles detected in this study were approximately 40 microns. While this was smaller that the previously described 50 micron diameter for VLPs (19, 26, 41), it is with in the range of native viral particles detected in oral wart lesions (11) as well as native HPV virus produced in organotypic cultures (8, 54). These particles were also detected in the media following treatment with NaB, and appeared identical to particles generated from cells not treated with the inducing agent (Figure 7C).

## Discussion

The association of oncogenic HPV16 with development of oropharyngeal cancer is well documented and HPV positive oral malignancies continue to rise in the general population (5, 17, 39). However, HPV16’s role in head and neck cancer is understudied compared to what is known about its role in cervical cancer. This paper describes the development and characterization of a simple cell culture system to study the HPV life cycle in differentiated oral keratinocytes. Here we show, in the presence and absence of a differentation agent, that HPV16 changed the differentation status of these oral keratinocytes, perhaps optimizing the cells for virus production. In this oral epithelial system viral genes were expressed from an oral cancer derived HPV16 isolate, gene products were functional, and viral particles were produced.

Determination of cell lines appropriate for studying the entire life cycle of HPV16 in cultured cells has made the study of HPV difficult. The primary challenge in the establishment of a culture system for HPV16 has been the tight link between virus maturation and keratinocyte differentiation (59). The use of primary keratinocytes grown as organotypic cultures that recapitulate the differentiation of epithelia has provided a system for investigating HPV16 virus production (34). Alternatively, in primary keratinocytes, induction of keratinocyte differentiation by either addition of CaCl_2_ or histone deacetylase inhibitors to the growth media or culturing in methylcellulose resulted in induction viral late gene expression (6, 35, 43). While each of these systems have provided major insights into viral gene expression and maturation, they require the isolation and maintenence of primary keratinocytes from human tissues and biopsies. Introduction of episomes into 293T cells followed by ectopic expression of L1 from an expression plasmid has also been used to produce infectious HPV16 (40). This system is dependent on the expression of ”codon optimized” L1 and L2 from expression plasmids for production of viral particles and packaging of episomes. The use of a Cre/lox system for the generation of HPV16 epsiomes was first reported by Lee et al. (32). A recombinant adenoviral vector cotaining the entire HPV16 genome flanked by loxp sequences was used to generate episomes in primary cervical keratinocytes following transduction with a adenovirus expressing Cre recombinase. A plasmid-based Cre/loxp system was subsequently developed and used to introduce HPV18 episomes into primary human foreskin keratinocytes (HFK) (54). Transfected cells were placed in organotypic culture to recapitulate epithelial differentation and produce HPV18 viral particles. To probe the full HPV lifecycle, the Cre-based HFK system required a continuous source of tissue for the generation of primary cell cultures as well as the use of raft cultures. HPV31 episomes have been introduced into monolayers of primary HFK cells (12). Treatment with CaCl_2_ (55) or growth in methylcellulose (12, 43) induced keratinocyte terminal differentation and resulted in late viral gene expression (E1^E4 transcripts) but the generation of infectious virus was not determined.

The cell line used in this study for investigation of the HPV life cycle was initially isolated from the oral cavity and immortalized with telomerase (7). Differentiated oral epithelial cells (OKF6tert1) cells demonstrated loss of heterogenetiy at the p16 locus, but retained normal growth and differentiation ability (7). These cells differentiated in organotypic rafts demonstrating K13 expression, a suprabasal marker specific to oral epithelial cells from non keratinizing oral mucosa (38). In the absence of NaB treatment or of HPV, the monomeric form of involucrin was consistently detected in OKF6tert1 cells. Cells were grown in serum free media where EGF and other growth factors were removed such that growth factors were not driving changes in differentiation (see methods section). The constitutive expression of monomeric involucrin, high levels of suprabasal cytokeratin, K10, and low levels of loricrin in this cell line, suggests that the cells were differentiated(Figure 1, Figure 8 A). In this state, OKF6tert1 cells were HPV replication competent and demonstrated a 1 log increase in viral DNA over a 72 hour time period (Figure 2). HPV16 expressing cells increased the terminal differentiation marker loricrin and decreased suprabasal and basal markers K10 and pser10H3, respectively; perhaps, achieving a state optimal for HPV replication. OKF6tert1 cells when exposed to differentiation agents, CaCl2 or NaB, were further differentiated moving to full terminal differentiation (Figure 1).

**Figure 8.**
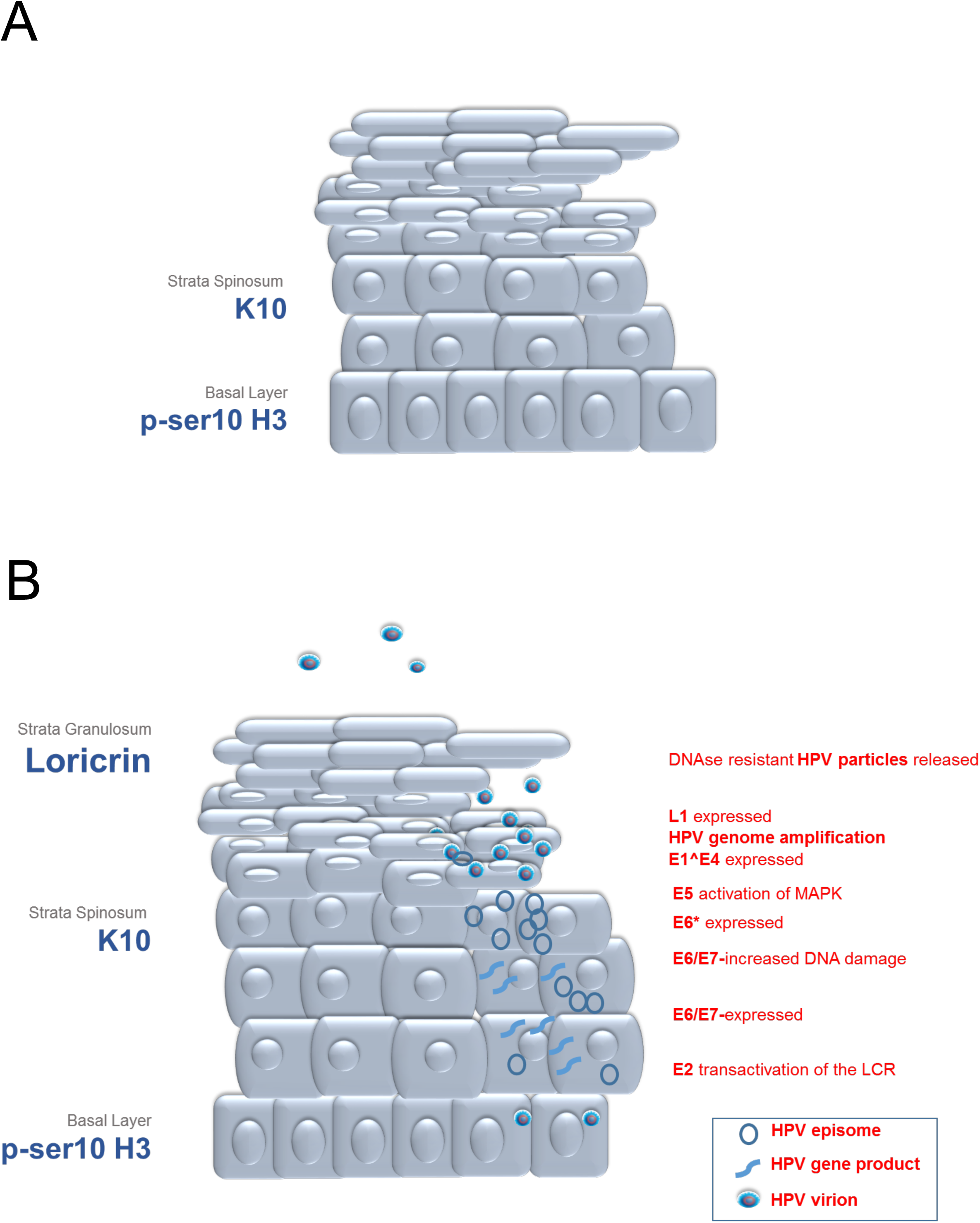
Model of HPV16 life cycle in transfected OKF6tert1 oral epithelial cells. **A.** A graphical representation OKF6 cells and associated differentiation markers. **B.** A graphical representation HPV infected oral epithelial cells. Viral amplification and release shown within the oral epithelium (center). Expressed epithelial differentiation markers are shown adjacent to epithelial layers (left). HPV gene expression and life cycle related cellular processes are shown adjacent to epithelial layers. HPV gene expression was tightly tied to movement toward terminal differentiation in the OKF-6 system (right).

HPV+ cervical carcinoma cells under go growth arrest after treatment with HDAC inhibitors including NaB (13). Detection of increasing loricrin and high molecular weight anti-involucrin immunoreactive proteins after treatment with NaB or CaCl2 in concert with decreasing suprabasal K10 indicated that full terminal differentation was induced in these cells (Figure 1). In the presence of increasing HDAC inhibition, HPV16 promoted a suprabasal phenotype while simultaneously suppressing terminal differentation (Figure 3). p53 inactivation in human keratinocytes causes squamous differentiation (14). The E6 viral oncogene inactivates p53 thus promotes differentiation. E6 mediated promotion of differentiation was consistent with increasedloricrindetection and decreased detection of both K10 and of the mitotic marker pS10H3 in oral cells expressing HPV16 compared to those expressing svGFP (Figure 1 and 3). Similar to what has been shown in HFK cells expressing HPV31, HPV16 alone appeared to decrease K10 expression levels (31). Likewise in CIN612 cells, that contained HPV31 episomes, increasing levels of both suprabasal and of terminal differntiation markers have been demonstrated (Langsfeld, 2016). However, in the presence of HPV16+ NaB,, K10 and pS10H3 levels increased and loricrin decreased but not to the same levels as detected in HPV16 negative cells (Figure 3). Thus it appears that HPV expressing cells in the presence of a differentiating agent modify the normal differentiation process. NaB mediated increases in terminal differentiation was associated with increased expression of E6/E7, E6* and E1^E4. However, changes in L1 were not detected, suggesting that baseline differentiation levels provided sufficient host factors for L1 splicing and expression. In sum, HPV expressing OKF6tert1 cells may be optimal for the HPV viral life cycle with a combined enhanced suprabasal phenotype, critical to early gene expression, and an enhanced terminally differentiated phenotype important to viral amplification, late gene expression and assembly (Figure 8 B, Table 1).

While other systems have generated important information on all aspects of the HPV viral life cycle, they have largely relied on previously laboratory adapted cloned HPV16 genomes. The system described here used DNA cloned directly from clinical isolates obtained from oral cancer samples as a source for generating HPV16 epsiomes in an oral keratinocyte cell line. The ability to PCR amplifiy whole genomes of HPV16 obtained from oral cancer biopsies provided a source of clinical isolates for the production of HPV16. The HPV16 episomes generated after introduction of these clinical HPV16 isolates into immmortalized oral keratinocytes were capable of directing viral gene expression. This was evident by the production of early viral transcripts encoding E6/E7, the production of late spliced transcripts (E1^E4, E6*, and L1), and the detection of viral proteins E7 and L1; all produced in the context of a differentiated oral keratinocyte (summarized in Table 1). Differentiated oral epithelial cells harboring HPV16 and/or its associated gene products demonstrated induction of the DNA damage response by E6/E7, activation of proliferative pathways by E5, transactivation of the HPV LCR by E2, and splicing of the E6/E7, E1^E4, and L1 transcripts - all hallmarks of HPV infection. The differentiated nature of these keratinocytes allowed for detection of L1 transcripts and L1 protein as determined by IFA, however, full terminal differentiation with NaB, did not provided any additional advantage to L1 expression. Interestingly, NaB mediated full terminal differentiation resulted in increased levels E1^E4 but decreased levels of L1 suggesting that the requirements for differentiation associated host factors were distinct for these viral late gene products. Transmission electron microscopy consistently detected a particle size of 40nM regardless of differentiation state. This size was considerably smaller than the 50-55 nM virus like particle (VLP) size (19, 26, 41). However, the size of particles detected in this report agreed with the sizes of native condensed viral particles detected in oral wart biopsies (11) and in organotypic cultures (8, 54). Thus, HPV16 episomes derived from oral cancer biopsied clinical isolates were capable of directing the full compliment of viral genes necessary for virus production in OKF6tert1 cells.

In sum, the OKF6tert1/HPV system provided several advantages for the study of the entire HPV16 life cycle, in that the cells were immortalized, easily cultured, and could be further differentiated (summarized in Table 1 A and B; Figure 8A). In this system, HPV influenced cell differentiation, perhaps creating an optimal situation for viral replication. Furthermore, the addition of NaB to this system recapitulated differentiation dependent viral gene expression. (Figure 8B). The utility of this system can be realized not only through the study of HPV pathogenesis but also through the testing of HPV-specific theraputics in oral epithelial cells.

## Acknowledgements

We would like to thank James G. Rheinwald for providing us with the OKF6tert1 cells used in this study and Marian Couch and Xiao Ying Yin for providing us with oral cancer biopsy DNA used to make whole HPV genomes. We also thank Scott Williams for providing reagents. This research was supported by NCI supplemental funding for HIV-associated malignancy research to the UNC Lineberger Cancer Center and the UNC CFAR (P30-CA016086), NIH/NIDCR and NIAID (OHARA) 1U01AI068636 and NIH/NIDCR 1R56DE023940-01.

