## Supplemental Table 1 for "Differentiated Oral Epithelial Cells Support the HPV Life Cycle"

Supplemental Table 1. Sequence of oligonucleotide primers used for PCR and qPCR

| Primer | Sequence |
| --- | --- |
| 16loxpF | 5’GCG*CCCGGG*ATAACTTCGTATAGCATACATTATACGAAGTTATGTGTATGTGTTTTTAAATGCTTGTGT3’ |
| 16loxpR | 5’GCG*AAGCTT*ATAACTTCGTATAATGTATGCTATACGAAGTTATACGTGTTTATTATACCATACATACAAACAC3’ |
| HPV16LCRREPF | 5’GCGACGCGTGACCTAGATCAGTTTCCTTTAGGACG3’ |
| HPV16LCRREPR | 5’GCGAGATCTATAAAATGTCTGCTTTTATACTAACCGGT3’ |
| HPV16E6EXF | 5’ CGCAAGCTTGGATGCACCAAAAGAGAACTGCA3’ |
| HPV16E6EXR | 5’ CGCGAATTCTTACAGCTGGGTTTCTCTACG3’ |
| HPV16E7EXF | 5’CGCAAGCTTATGCATGGAGATACACCTACATTG3’ |
| HPV16E7EXR | 5’ CGCGAATTCTTATGGTTTCTGAGAACAGATGGG3’ |
| HPV16E2EXPF | 5’ GGGAAGCTTGCCATGGAGACTCTTTGCCAACGTTT3’ |
| HPV16E2EXPR | 5’GGGACTAGTTCATATAGACATAAATCCAGTAGACACTGT3’ |
| HPV16E1^E4F | 5’CCCCATCTGTTCTCAGAAACC3’ |
| HPV16E1^E4R | 5’GGCCAATGTCTGCCTAATAA3’ |
| HPV16L1splcF | 5’ACATGTCTCCAATCCTCACTGCAT3’ |
| HPV16L1splcR | 5’GCGTGCAACATATTCATCCGTGCT |
| HPV16JNCF | 5’CCAAAATTTACATTAGGAAAACGAA3’ |
| HPV16JNCR | 5’TTTATGTTGCATGACACAATAGTT3’ |
| GAPDHF | 5’GAAGGTGAAGGTCGTAGTC3’ |
| GAPDHR | 5’GAAGATGGTGATGGGATTTC3’ |
